# Recombination alters the receptor binding and furin cleavage site in novel bat-borne HKU5-CoV-2 coronavirus

**DOI:** 10.1101/2025.04.17.649446

**Authors:** Ting-Yu Yeh, Vincent Tsai, Samuel M. Liao, Chia-En Hong, Feng-Yu Kuo, Yen Chun Wang, Michael C. Feehley, Patrick J. Feehley, Yi-Chen Lai, Gregory P. Contreras

**Affiliations:** Auxergen Inc., Rita Rossi Colwell Center, Baltimore, Maryland, MD 21202, USA; Auxergen S.r.l., Tecnopolis Science and Technology Park of the University of Bari, University of Bari, Valenzano BA, Italy; Taipei Wego Private Senior High School, Beitou District, Taipei City, Taiwan; Davis Senior High School, Davis, CA, USA; Henry M. Gunn High School, Palo Alto, CA, USA; Taipei Municipal Chien Kuo High School, Taipei City, Taiwan; Taipei Municipal Chenggong High School, Taipei City, Taiwan; Department of Biophysics, Johns Hopkins University, Baltimore, MD 21218, USA; Department of Radiology, Taipei Veterans General Hospital, Taipei city, Taiwan; School of Medicine, National Yang Ming Chiao Tung University, Taipei City, Taiwan

## Abstract

HKU5-CoV-2 is a new bat-infecting coronavirus phylogenetically related to MERS-CoV. It has recently been confirmed that HKU5-CoV-2 can enter human cells and organoids *in vitro* via the ACE2 receptor, raising concerns about its pandemic potential due to zoonotic spillover. Whether recombination has an influence on HKU5-CoV-2 is completely unclear to date. Here we report the first evidence of HKU5-CoV-2 viral recombination, in association with mutations at the receptor binding domain (RBD) and S1/S2 furin cleavage site (FCS) of the spike protein. Using linkage disequilibrium and haploblock analysis, we identified that 167 recombination hotspots and 27 haploblocks.

SNP23016/23043/23064/23156/23193/23285 at the RBD and SNP23833/23847 at the FCS are recombinant hotspots. Our results suggest that recombination may lead to the substitution at RBD residue 498 (Thr498Val/Val498Thr, Thr498Ile/Ile498Thr), which Thr498 directly contacts the ACE2 receptor.

Recombination also causes Ser723 deletion/insertion and Ser729Ala substitution at the FCS. These mutations could affect host tropism and change furin cleavage activity. Our results indicate that recombination has played a critical role in HKU5-CoV-2 evolution and infectivity.

## Introduction

HKU5-CoV-2 merbecovirus is a newly discovered coronavirus from *Pipistrellus* spp. bats in China. Phylogenetically related to MERS-CoV, HKU5-CoV-2 can enter host cells via the ACE2 receptor present in many birds and mammals, including humans. ^1^ Although there is no evidence of human transmission to date, HKU5-CoV-2 poses a risk of zoonotic spillover with pandemic potential. It is well known that recombination quickly shapes the pathogenesis and evolution of coronaviruses like SARS-CoV-2, even within 3 weeks of the human transmission.^2^ However, whether recombination has influenced HKU5-CoV-2 is completely unknown. Here we report the first evidence that recombination drove HKU5-CoV-2 diversity and has led to alterations in the receptor binding domain and S1/S2 furin cleavage site.

## Results and Discussion

Six HKU5-CoV-2 sequences are available in GenBase (C-AA08189 to C-AA08194).^1^ Linkage disequilibrium (LD) analysis of 765 single nucleotide polymorphisms (SNPs) shows the nonrandom feature of LD and recombination (Figure 1A and 1B), with 167 recombination hotspots and 27 haploblocks (Figure 1C). SNP23016/23043/23064/23156/23193/23285 in the receptor binding domain (RBD) of the spike protein are hotspots for intradomain or long-range recombination (Figure 2A-2C, 2F). Recombination occurred between SNP23155/23156, an event that may cause Thr498Val/Val498Thr or Thr498Ile/Ile498Thr substitutions. Thr498 directly contacts the human ACE2 receptor Asp30/His34/Glu37, and RBD Gly499/Asn503, which are also receptor binding residues (Figure 2D-2E).^1^ Substitutions at the RBD residue 498 suggests that recombination can change receptor binding, host range, and/or viral infectivity of HKU5-CoV-2.

**Figure 1.**
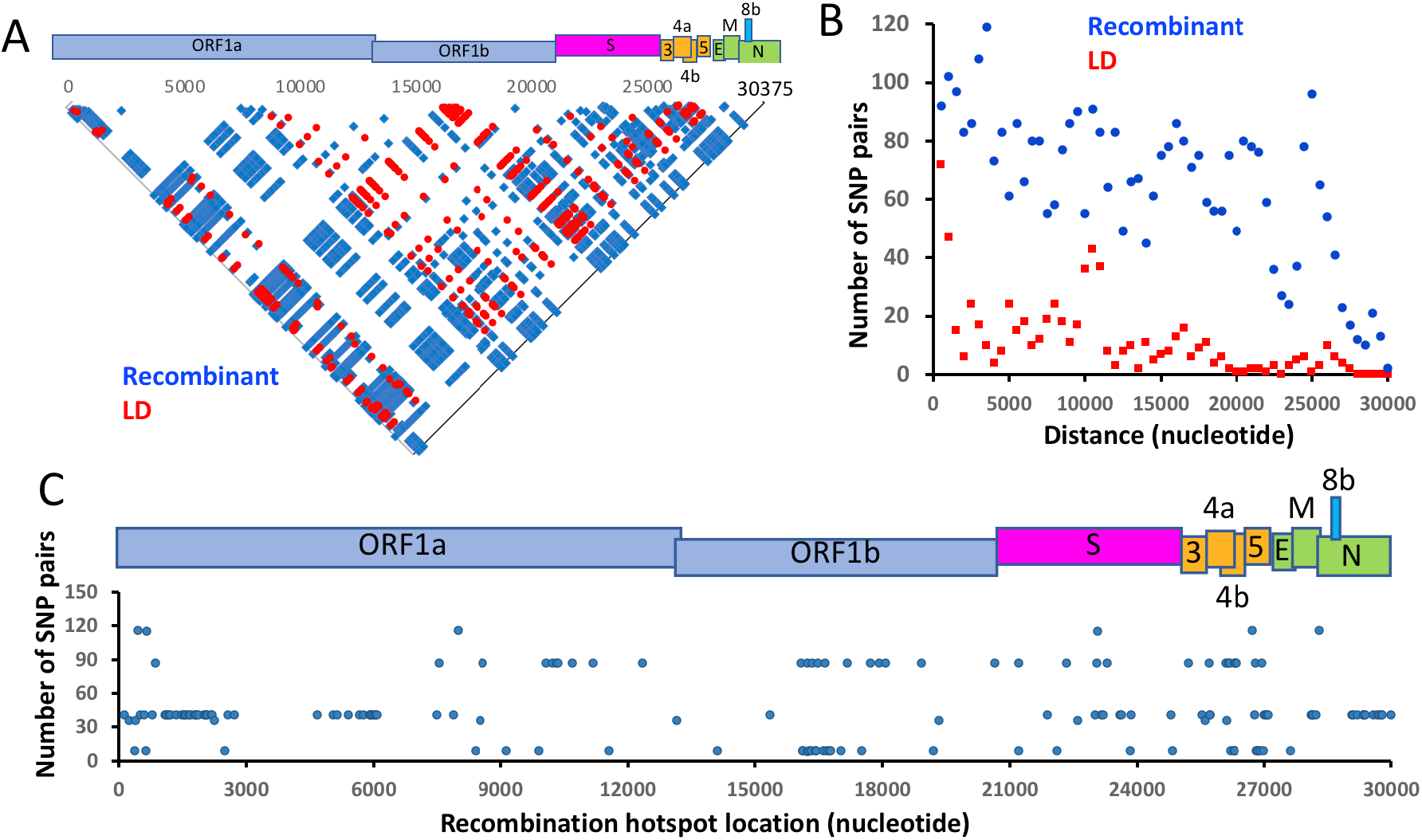
LD analysis of HKU5-CoV-2 genome. (A) Pairwise plots of LD (N=666, red) and recombinant (N=3896, blue) SNP pairs. LD is defined if pairs for which the value of logarithm of the odds (LOD) >2, the squared coefficient of correlation (*r*^2^) =1, and the high value of 95% confidence interval (CIhigh) bounds for *D*’ =1. Recombination is defined if pairs for which the upper CI bound of *D*’ is below 0.9. (B) The nonrandom distribution of LD (red) and recombinant (blue) SNP pairs are shown by plotting the pair numbers (Y-axis) against the genomic distance between that pair (x-axis). (C) The locations of HKU5-CoV-2 recombination hotspots, which the numbers of their associated recombinant SNP pairs are shown in Y-axis. Genomic organization of HKU5-CoV-2 is shown above (A and C).

**Figure 2.**
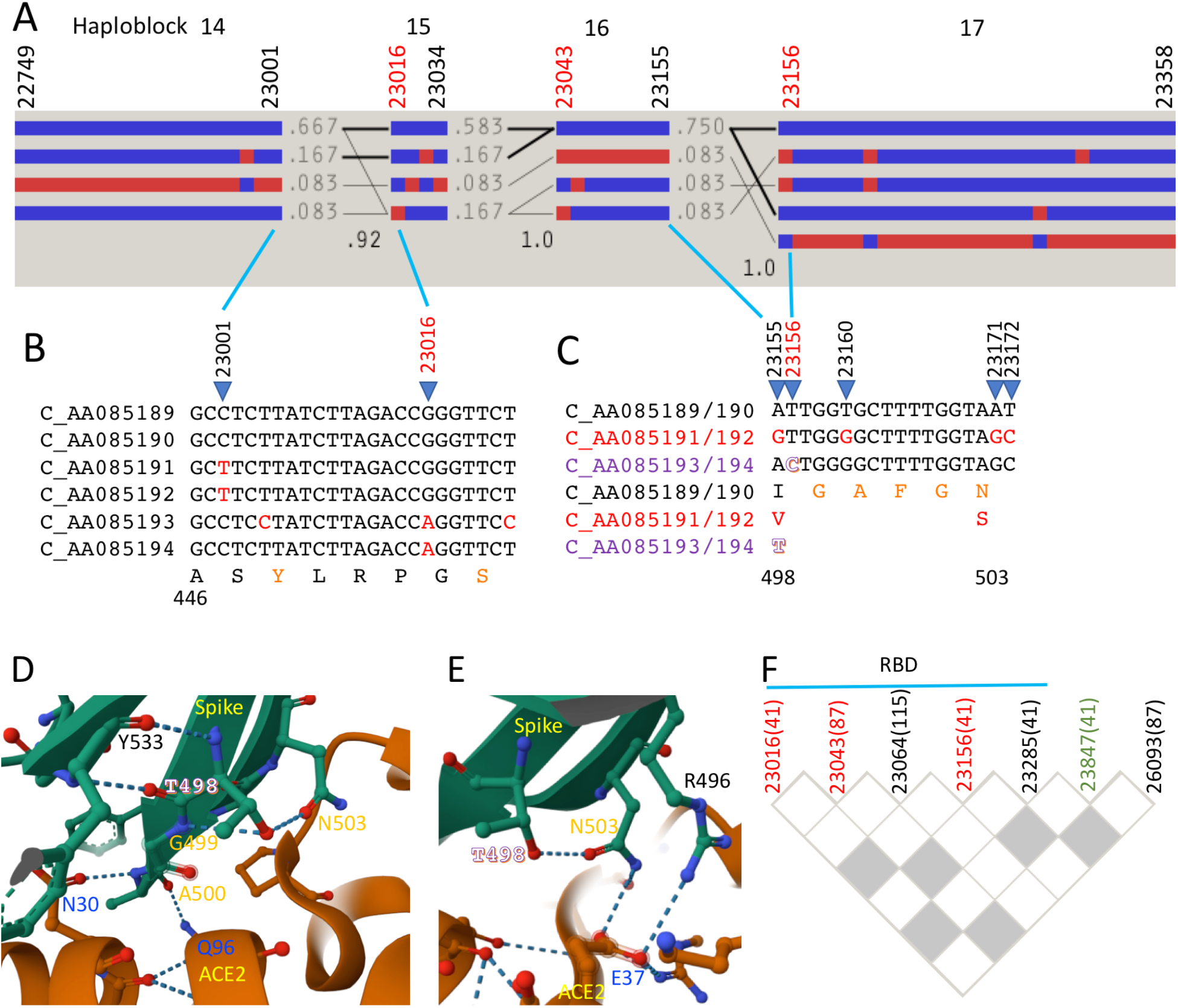
Recombination of HKU5-CoV-2 RBD. (A) Haplotype block (14 to 17) organization of HKU5-CoV-2 RBD sequences. The Hedrick’s multi-allelic *D*’ statistics in the crossing areas represents the recombination level between two haploblocks. Haplotype frequencies are shown as the numbers next to each haplotype block, or > 10% (thick lines) or > 1% (thin lines). (B and C) Alignment of the adjacent sequences of recombinant breakpoints of SNP23016 (B) and SNP23156 (C). ACE2 receptor binding residues^1^ are colored in orange. T498 is outlined (purple). (D and E) Protein structure around RBD (green) T498 binding to the ACE2 receptor (brown) is illustrated by RCSB (PDB: 9JJ6)^1^. RBD receptor binding sites: G499, A500, N503 (orange). ACE2 receptor’s spike binding sites: N30, E37, Q96 (blue). Hydrogen bonds are shown by dashed lines. (F) Pairwise LD plots of recombinant SNP pairs (white squares) in RBD, FCS (SNP23847, green), or the intergenic region between spike and E protein (SNP26093). The numbers of recombinant SNP pairs are associated with each SNP. The breakpoints in haploblock 15, 16, and 17 (A) are colored in red.

It has been suggested that the S1/S2 furin cleavage site (FCS) appeared independently multiple times in the evolution of the coronavirus spike protein.^3^ We find that the HKU5-CoV-2 FCS contains several mutations (Figure 3A). Haploblock analysis revealed that recombination occurred at SNP23830/23833 (Figure 3B).

**Figure 3.**
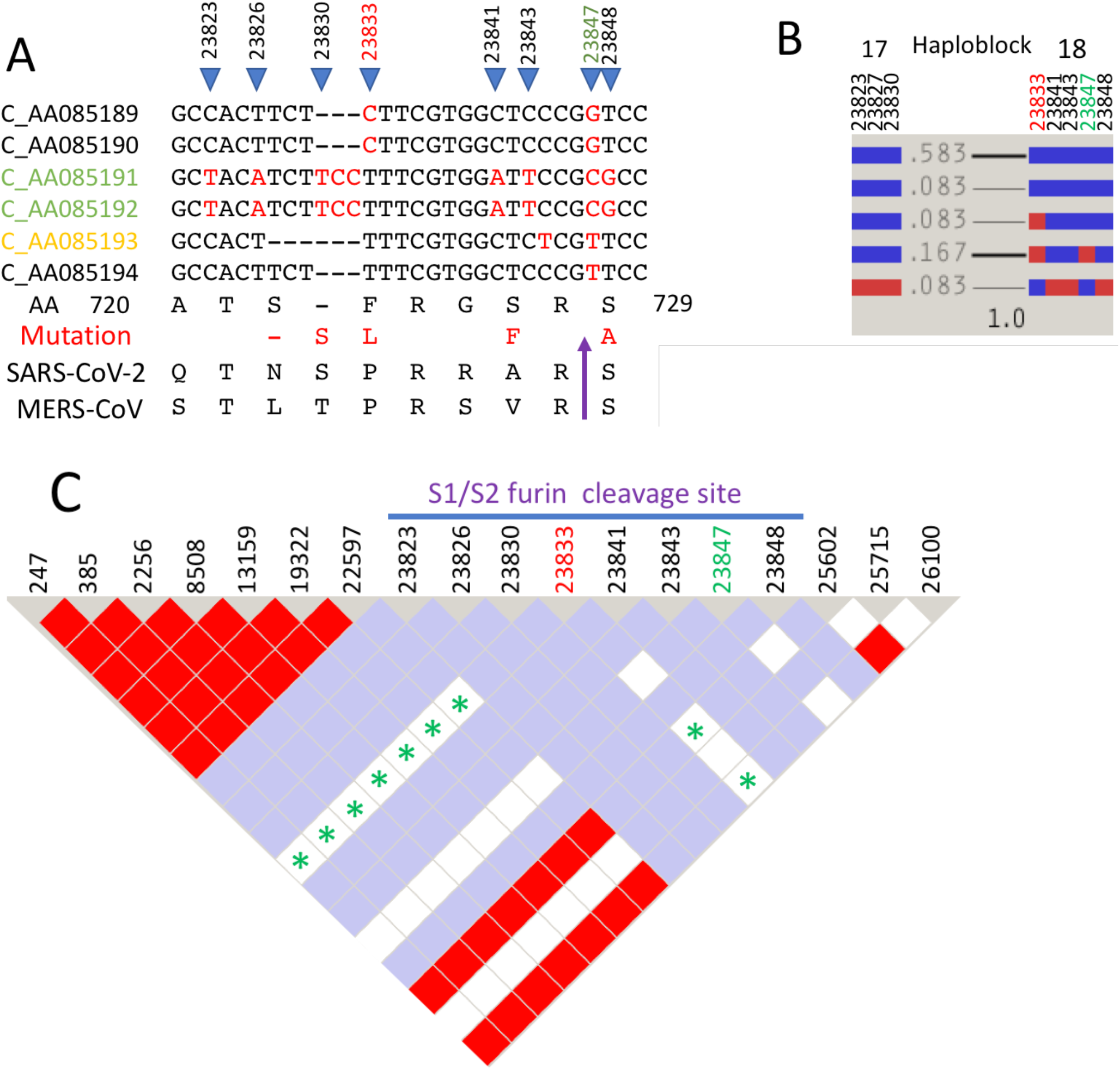
Recombination of HKU5-CoV-2 FCS. (A) Alignment of FCS sequences. Furin cleavage site (purple arrow), sequences with Ser272/273 deletion (yellow), or S729A substitution (green) are colored. Recombination hotspots SNP23833 and SNP23847 are labeled in red. (B) Haplotype block organization of (A). (C) HaploView LD map of FCS and other SNPs. Red squares indicate high levels of LD, and white and blue squares represent low levels of LD. Recombinant SNP pairs are marked with asterisks.

Moreover, SNP23833 and SNP23847 also form 9 and 41 recombinant SNP pairs, respectively, indicating that the FCS has been a crossover hotspot (Figure 3C and 2F). Additionally, the FCS is often lost during virus adaptation to cell culture by either deletions or point mutations.^4^ Taken together, these results suggest that recombination may cause Ser723 deletion/insertion and Ser729Ala substitution, both of which could change furin cleavage activity (e.g. C_AA085189/190/194 versus C_AA085191/192/193).

Another non-LD algorithm, Recco, identified 10 putative recombination breakpoints in NSP4, NSP13, NSP14, and the transmembrane domain of the spike protein (Table 1). Other breakpoints (SNP18065/19205/25521) were also confirmed by LD and haploblock analysis. It is reasonable to speculate that recombination of HUK5-CoV-2 virus is underestimated. Additionally, it has also been reported that recombination events can alter the host tropism of MERSr-COVs and BtCov-422 due to broader receptor usage.^5^ Therefore, RBD and FCS variations caused by recombination and mutations could significantly affect viral entry and infectivity. Genomic surveillance of HUK5-CoV-2 and its recombinants in wild bat populations could be the key strategy to mitigate zoonotic transmission and future outbreaks.

**Table 1.** Recco analysis of HKU5-CoV-2 sequences*. The recombination breakpoints are verified by LD (recombinant SNP pairs^#^) and haploblock analysis (between or at the end of haploblocks).

| **Recombinant fragment (nucleotide)\*** | **Mutation cost\*** | **SNP (recombinant SNP pair numbers)<sup>#</sup>** | **Location** | **Haploblock** |
| --- | --- | --- | --- | --- |
| 8893-8997 | 381.2<br>($p=0.04$) | 8892 (0) | Polyprotein 1ab, AA 2879-2913, NSP4 transmembrane domain | 10 |
| 18066-18397 | 410.2<br>($p<10^{-3}$) | 18065 (86) | Polyprotein 1ab, AA 5937-6046, NSP13 Helicase, NSP14 ExoN domain | 12 |
| 19206-19292 | 410.2<br>($p<10^{-3}$) | 19205 (9) | Polyprotein 1ab, AA 6317-6345, NSP14, N7-MTase active site | 12 (end) |
| 25522-25563 | 394.2<br>($p=0.017$) | 25521 (41) | spike protein AA 1286-1299, transmembrane domain | 19, 20 |

## Materials and Methods

Genomic sequences of HKU-CoV-2 (BtHKU5-CoV-2-153, BtHKU5-CoV-2-155, BtHKU5-CoV-2-023, BtHKU5-CoV-2-028, BtHKU5-CoV-2-381, and BtHKU5-CoV-2-441) were obtained from Genbase database (National Genomics Data Center, China, https://ngdc.cncb.ac.cn/genbase/?lang=en). Accession numbers: C_AA085189.1, C_AA085190.1, C_AA085191.1, C_AA085192.1, C_AA085193.1, and C_AA085194.1. The nucleotide and SNP number are based on the C_AA085191.1 sequence in this study. Sequence alignment and linkage disequilibrium (LD) analysis using HaploView (https://www.broadinstitute.org/haploview/haploview) are previously described. ^6,7^ Only mutations shown in the same allele in more than two sequences, but not the singletons, were considered SNPs and analyzed in this study. The *D*’ value was calculated by normalizing the degree of nonrandom association between two alleles (*D*), with its maximum possible value *D*_max_. Haplotype blocks with recombination measurements were determined with the solid spine of LD. ^7^

The viral sequences were analyzed by Recco program, which scores the cost of obtaining one of the sequences from the others by mutation and recombination.^8^ Recombinant fragments and breakpoints were measured by parameter α, weighting recombination cost against mutation cost. Only the fragments with significant mutation cost (>5) and *p* value <0.05 were considered as recombinant fragments.

## Declaration of interest

TYY is the CSO of Auxergen Inc and Auxergen s.r.l.. GPC is the CEO of Auxergen Inc and Auxergen s.r.l.. Other authors have no conflict of financial interest.

## Funding

No funding

## Ethical approval

None declared.

## Author Contributions

Yeh and Contreras had full access to all the data in the study and take responsibility for the integrity of the data and the accuracy of the data analysis.

Concept and design: Yeh

Acquisition, analysis, or interpretation of data: Yeh, Tsai, Laio, Hong, Kuo, Wang, Feehley MC, Feehley PJ, Lai

Drafting of the manuscript: Yeh, Contreras

Critical revision of the manuscript for important intellectual content: Yeh, Contreras

Statistical analysis: Yeh, Tsai, Laio, Hong, Kuo, Wang, Feehley MC, Feehley PJ, Lai

Supervision: Yeh, Contreras

## Notes

### Summary of Updates

New Figure 1: LD analysis and location of recombination hotspots. New Figure 2C: RBD Thr498 as the recombination hotspot with Val and Ile substitution. New Figure 2D and 2E: Protein structure of RBD Thr498 with ACE2 receptor.

